# Study on genetic differentiation of *Schistosome japonicum* intermediate hosts *Oncomelania hupensis robertsoni* in hilly regions of China: using the complete mitochondrial genome

**DOI:** 10.1101/2023.11.05.565742

**Authors:** Jing Song, Hongqiong Wang, Shizhu Li, Zongya Zhang, Chunying Li, Jihua Zhou, Meifen Shen, Peijun Qian, Wenya Wang, Yun Zhang, Chunqiong Chen, Lifang Wang, Jiayu Sun, Yuwan Hao, Chunhong Du, Yi Dong

## Abstract

**Objective:** *Oncomelania hupensis robertsoni* is the only intermediate host of *Schistosoma japonicum* in western China, its genetic differentiation directly impacts the susceptibility of *Schistosoma japonicum.* This study aimed to sequence the complete mitochondrial genome of *Oncomelania hupensis robertsoni* Yunnan strain and analyze the genetic differentiation of *Oncomelania hupensis robertsoni* in hilly regions of China.

**Methods:** Samples were from 14 administrative villages in Yunnan Province of China, with 30 *Oncomelania hupensis* per village, and the complete mitochondrial genome was sequenced. Additional, we retrieved 14 other region *Oncomelania hupensis* of complete mitochondrial sequences from GenBank, and a comprehensive analysis of the genetic differentiation of *Oncomelania hupensis robertsoni* was conducted by constructing phylogenetic trees, calculating genetic distances, and analyzing homogeneity.

**Results:** A total of 26 complete mitochondrial sequences were determined. The length of genome ranged from 15,181 to 15,187 bp, and the base composition of the genome was A+T (67.5%) and G+C content (32.5%). This genome encoded 37 genes, including 13 protein-coding genes, 2 rRNA genes, 22 tRNA genes and a non-coding region rich in A+T. Using the Philippines genotypes as outgroup, the phylogenetic trees and homology analysis confirmed the existence of two distinct phylogroups, *Oncomelania hupensis robertsoni* and the remaining 9 provincial genotypes. *Oncomelania hupensis robertsoni* is subdivided into *Oncomelania hupensis robertsoni* Yunnan strain and Sichuan strain, with a genetic distance of 0.0834. *Oncomelania hupensis robertsoni* Yunnan strain is subdivided into two subbranches, “Yunnan North” and “Yunnan South”, with a genetic distance of 0.0216, and the samples exhibited over 97% homology.

**Conclusion:** *Oncomelania hupensis robertsoni* Yunnan strain exhibits a higher level of genetic homology and clear north-south differentiation, the distribution characteristics were closely associated with watershed distribution. This work reported the first mitochondrial genome of *Oncomelania hupensis robertsoni* Yunnan strain, which could be used as an important reference genome for *Oncomelania hupensis*, and also provide a theoretical basis for explaining the distribution pattern of *Oncomelania hupensis robertsoni* and control of schistosomiasis.

**Author Summary:** *Oncomelania hupensis* (*O. hupensis*) is the only intermediate host of *Schistosoma japonicum* (*S. japonicum)*, *O. hupensis* residing in different geographical regions display morphological differences and genetic variations, along with varying susceptibility to *S. japonicum*. In this study, we sequenced 26 complete mitochondrial genome of *O. hupensis robertsoni* Yunnan strain (*O. h. r.* Yunnan strain), the length of genome ranged from 15,181 to 15,187 bp, and the base composition of the genome was A+T (67.5%) and G+C content (32.5%). This genome encoded 37 genes, including 13 protein-coding genes, 2 rRNA genes, 22 tRNA genes and a non-coding region rich in A+T. Additional, we retrieved 14 other region *O. hupensis* of complete mitochondrial sequences from GenBank. The phylogenetic trees and homology analysis confirmed that *O. hupensis robertsoni* is subdivided into Yunnan strain and Sichuan strain, and *O. h. r.* Yunnan strain is subdivided into two subbranches, “Yunnan North” and “Yunnan South”, the samples exhibited over 97% homology. This work reported the first mitochondrial genome of *O. h. r.* Yunnan strain, which could be used as an important reference genome for *O. hupensis*, and also provide a molecular biology-based theoretical foundation for understanding the genetic differentiation of *O. hupensis*.

## 1 Introduction

*Oncomelania hupensis* (*O. hupensis*) is the only intermediate host of *Schistosoma japonicum* (*S. japonicum)*, and its distribution area directly determines the epidemic range of schistosomiasis[1]. Despite differences in morphology among *O. hupensis* from different countries and regions, they can still be classified as the same species based on their genetic characteristics of reproductive isolation and chromosomal homology[2, 3]. While it’s currently recognized that there are eight subspecies of *O. hupensis*, the classification of these subspecies has been controversial due to factors such as identification methods and sample sources. Nevertheless, the classification of *O. hupensis robertsoni* (*O. h. robertsoni*) remains relatively well-defined. In China, *O. hupensis* primarily inhabits the 12 southern provinces in the middle and lower reaches of the Yangtze River[4], its geographical distribution range is extensive, with notable climate variations and complex array of environmental types[5]. *O. hupensis* residing in different geographical regions display morphological differences and genetic variations, along with varying susceptibility to *S. japonicum*[4, 6, 7]. Considering the close genetic interaction between *S. japonicum* and its intermediate host, the *O. hupensis*, in terms of coevolution[8, 9], and the fact that *O. hupensis* with higher genetic diversity are more susceptible to *S. japonicum*[10], conducting research on the genetic differentiation and classification of *O. hupensis* is of great significance for understanding the transmission of schistosomiasis and guiding disease control measures.

In recent years, mitochondrial DNA (mtDNA) has found widespread application as a molecular marker in numerous biosystematics studies[11–14]. However, current research on *O. hupensis* mtDNA mostly focuses on individual gene fragments such as cytochrome c oxidase1 (COX1) and cytochrome b (CYTB)[5, 10, 15–19], the results obtained have certain limitations to reveal the *O. hupensis* of population structure and genetic variation[20, 21].

Based on the morphological characteristics, biological traits, and molecular markers, *O. h. robertsoni* can be further divided into the Yunnan strain and Sichuan strain[4]. Compared with other subspecies, the morphological characteristics of *O. h. robertsoni* are smooth shell surface without longitudinal ribs, no labial ridges, and uniform growth of each conch layer[18]. Population genetic experiments have revealed that Yunnan Province is the likely place of origin for *O. hupensis* in mainland China (30-31). After initially entering Yunnan province from the Himalayan Mountains, *O. hupensis* subsequently migrated to the Sichuan Plain, which is connected to the Yangtze River, and then dispersed further to the east coast of mainland China. Given the role of Yunnan Province in genetic differentiation of *O. hupensis* in mainland China, it is very important to study the genetic differentiation and classification of *O. h. robertsoni* Yunnan strain (*O. h. r.* Yunnan strain). However, only one study focused on the complete mitochondrial genome of *O. hupensis*[17], but lacking the Yunnan Province, this merits more research of mitochondrial genome of *O. h. r.* Yunnan strain attention. In addition, endemic areas for schistosomiasis in Yunnan Province exhibit a distinct “island-like” pattern[22], prompting many researchers to speculate about the potential existence of on unique genetic characteristics within this regional *O. hupensis*.

The study herein aimed to sequence the complete mitochondrial genome of the *O. h. r.* Yunnan strain, and to explore intraspecific developmental relationships of the *O. h. robertsoni* and the differentiation characteristics of *O. h. r.* Yunnan strain. Our results may provide a molecular biology-based theoretical foundation for understanding the genetic differentiation of *O. hupensis* and how population structure of intermediate host *O. hupensis* influences susceptibility to *S. japonicum* infections, and guide the strategies of schistosomiasis control.

## 2 Methods

### 2.1 Source of *O. hupensis*

In the existing snail distribution area of Yunnan Province, 14 administrative villages in 10 schistosomiasis-endemic counties of 3 cities were selected as snail collection sites after considering various factors such as snail density, geographical location, altitude, water system, and environmental type (**Figure 1** **Distribution of sampling points of *O. hupensis* in Yunnan Province of China)**. 14 administrative villages were Leqiu (NJ1) and Anding (NJ2) in Nanjian County, Caizhuang (MD1) in Midu County, Xiaoqiao (XY2) in Xiangyun County, Qiandian (EY1) and Yongle (EY2) in Eryuan County, Wuxing (DL2) and Xiaocen (DL3) in Dali City, Dianzhong (WS1) in Weishan County, Lianyi (HQ1) in Heqing County of Dali Prefecture, Yangwu Village (YS3) in Yongsheng County, Sanyi Village (GC1) and Dongyuan Village (GC2) in Gucheng District of Lijiang City, and Cangling Village in Chuxiong County of Chuxiong Prefecture (CX2). One field snail collection site was selected in each administrative village to collect *O. hupensis* samples.

**Figure 1.**
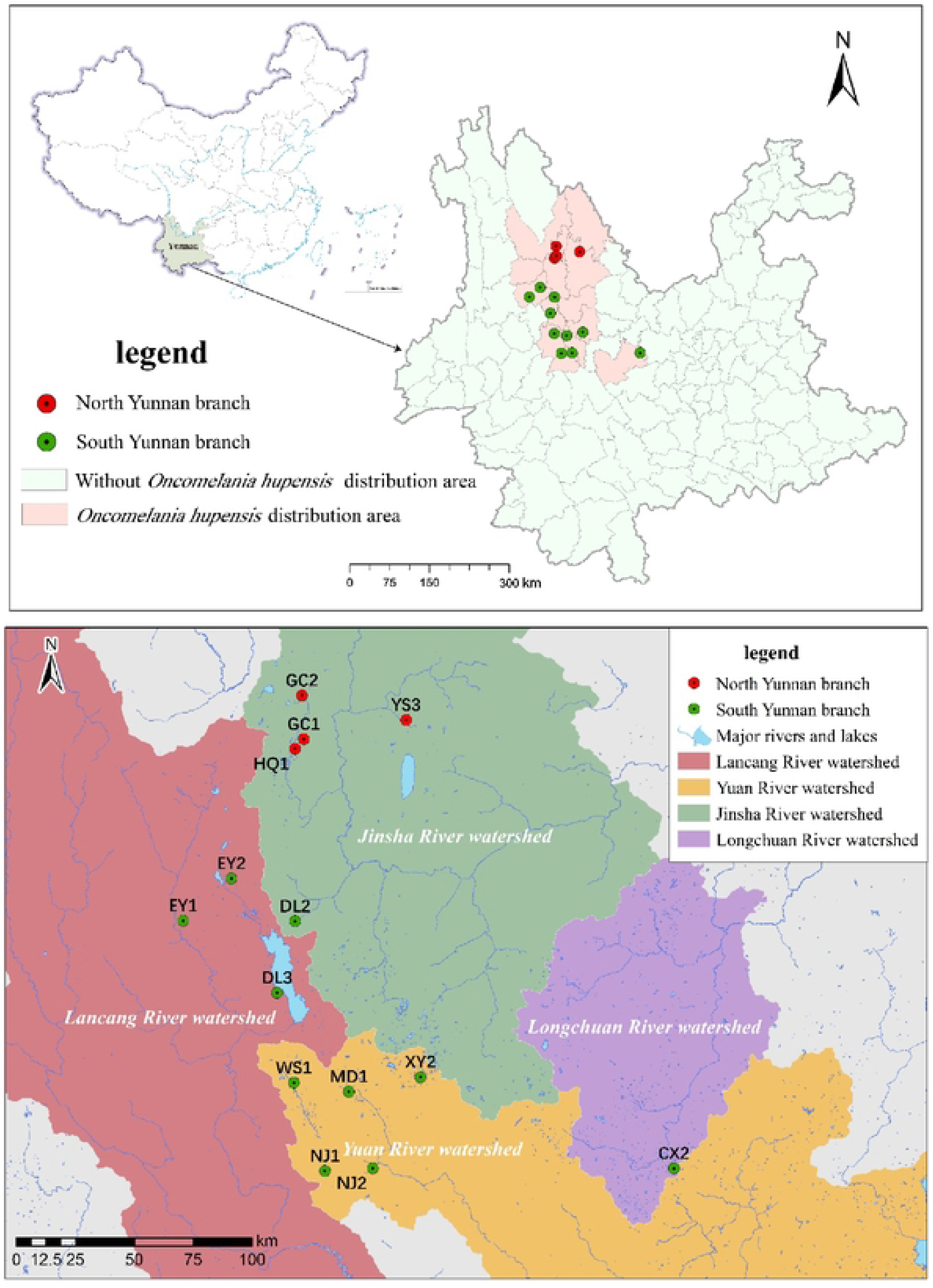
Distribution of sampling points of *O*. *hupensis* in Yunnan Province of China

Approximately 200 snails were randomly sampled from each point, and brought back to the laboratory to remove dead snails by crawling method, and identified whether the snails were infected with schistosomes by cercariae shedding method[23].

### 2.2 Extraction of total DNA from *O. hupensis*

The muscle tissue of the abdominal foot of a single *O. hupensis* was taken and the genomic DNA was extracted using the Qiagen extraction kit [Qiagen Enterprise Management (Shanghai) Co., Ltd].

### 2.3 Primer design and PCR amplification

The primers and PCR amplification reaction conditions were based on the literature[24] (**Table 1**). The entire genome was divided into 16 overlapping fragments according to the sequence length, and the primers were designed separately, except for the reverse primer of primer pair 10 and the primer pair 14, which were not consistent with the literature. The primer synthesis, PCR amplification, PCR product purification and sequence sequencing were performed by Shanghai Xianghong Biotechnology Co.

### 2.4 Sequence assembly and editing

All sequence fragments were filtered to remove those that were not normally reported, such as PCR “amplification failure” and sequencing “bimodal mutation”. The SeqMan module of the DNAstar software package was used to splice the 16 sequences of each sample, remove the sequences that could not be assembled into closed circular forms. The overlapping regions of adjacent sequences that could be assembled into closed circular forms were trimmed, and the entire sequence was manually checked for base identification based on peak shape. The assembled and edited sample sequences were submitted to GenBank for sequence alignment and direction confirmation. If the sequence orientations were inconsistent, the reverse complementary sequences of the sample sequences were obtained using the free online tool “SMS2 Nanjing Detai Biological Mirror”, and then submitted to GenBank for sequence alignment. Based on the gene order of the nearest mitochondrial sequence (the closest mitochondrial gene sequence to all sample sequences is the SCXC strain of *O. hupensis* mitochondrion, GenBank ID: JF284691), the sample sequences were trimmed and ordered to have the same starting gene sequence as the COX1 gene in the SCXC strain of O. hupensis mitochondrion.

### 2.5 Sequence alignment

The edited self-test sequences and all complete mitochondrial sequences of *O. hupensis* retrieved from GenBank were aligned using MEGA11.0.9 software.

### 2.6 Sequence composition analysis

The MEGA-formatted alignment results were imported into MEGA11.0.9 software, and the sequence component analysis was performed using the menu “statistics-Nucleotide Composition”. The start and end points of each gene were determined by combining MEGA11.0.9 software with NCBI’s BLAST to perform homology-based gene annotation analysis.

### 2.7 Construction of phylogenetic tree

Phylogenetic trees were constructed using MEGA11.0.9 software by the four methods of Maximum Parsimony (MP) (parameter settings: MP Search Method: SPR, No. of Initial Tree:10, MP Search level:1, Max No. of Tree to Rain:100), Maximum Likelihood (ML) (parameter settings: Model/Method: Tamura-Nei model, ML Heuristic Method: NNI, Initial Tree for ML: Make initial tree automatically), Minimum Evolution (ME) (parameter settings: Model/Method: Maximum Composite Likelihood, Substitutions to Include: d: Transitions+Transversions), and Neighbor-Joining (NJ) (parameter settings: Model/Method: Maximum Composite Likelihood, Substitutions to Include: d: Transitions+Transversions), respectively, and each method had repeats of 1000 bootstrap value .

### 2.8 Sequence genetic distance and homology analysis

Genetic distances within and between groups were calculated using the “distance” menu in MEGA11.0.9 software based on alignment results. Sequence homology was calculated using the “View-Sequence distance” option in MegAlign of the DNAstar package. The average nucleotide similarity among 40 snail sequences was calculated using Fastani software, homology analysis was performed by comparing genome sequence similarities combined with Blast, and the Average nucleotide identity (ANI) heat map was generated by Tbtools.

## 3 Results

### 3.1 Sequence assembly, editing, and alignment

After assembling and editing the 89 “normal report” sequences, 26 complete mitochondrial genome sequences from 13 sampling sites were obtained (the sequence from Dianzhong Village in Weishan County was not obtained successfully) and submitted to GenBank. The number of sequences obtained from each sampling sites ranged from 1 to 4.

In addition, 14 complete mitochondrial genome sequences of *O. hupensis* in this study were used in constructing phylogenetic trees, calculating genetic distances, and analyzing homogeneity with 14 other reference strains retrieved from GenBank, including those from the Philippines (JF284698.1 FLB), Sichuan Province (JF284697.1 SCMS, JF284691.1 SCXC), Fujian Province (JF284695.1 FJFQ), Zhejiang Province (JF284694.1 ZJJH), Guangdong Province (MN200239.1 Guangdong), Jiangsu Province (JF284688.1 JSYZ), Anhui Province (JF284686.1 AHGD-1, JF284687.1 AHGD-2), Hunan Province (JF284692.1 HNYY), Hubei Province (JF284689.1 HBGA, JF284690.1 HBJL), Jiangxi Province (JF284693.1 JXSR), and Guangxi Province (JF284696.1 GXBS).

### 3.2 Sequence base composition analysis

The length of the complete sequence of *O. h. r.* Yunnan strain mitochondrial genes was 15,181bp∼15,187bp, with an average length of 15,185bp, and the average contents of A, T, C and G bases were 29.7%, 37.8%, 15.6% and 16.9%, respectively, in which the A+T content (67.5%) was significantly higher than the G+C (32.5%) content. The complete mitochondrial genome of *O. h. r.* Yunnan strain comprised 37 genes, including 13 protein-coding genes, 2 rRNA genes, 22 tRNA genes, and a non-coding region rich in A+T. Detailed information about the organization of *O. h. r.* Yunnan strain mitochondrial genome in 13 sampling sites are presented **eTable1∼eTable9 in the supplement.**

For example, in NJ1-01 sampling site, there weas 19 intergenic regions totaling 272bp, ranging in length from 1bp to 68bp, with the largest intergenic region located between ND5 and trnF (gaa) (68bp), and 5 gene overlap regions totaling 17bp (**Figure 2** Representative genome map representing the mitochondrial genome circular molecule of *O. h. r.* Yunnan strain).

**Figure 2.**
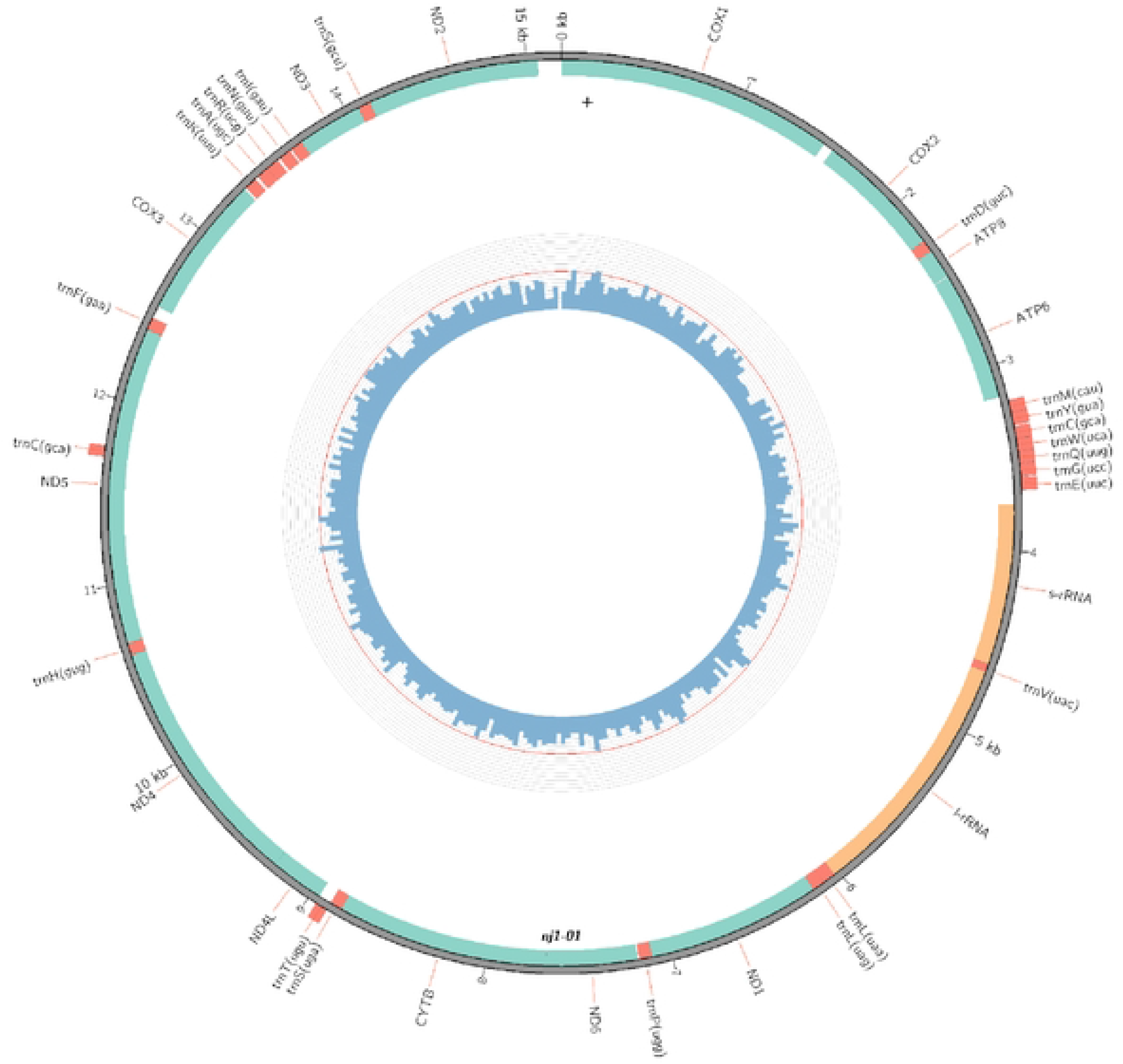
Representative genome map representing the mitochondrial genome circular molecule of *O. h. r.* Yunnan strain. The colored squares distributed inside and outside the circle represent different mitochondrial genes, gene taxa of the same function are represented using the same color.

### 3.3 Phylogenetic tree construction

The topology of the phylogenetic tree constructed by the four methods of MP, ML, ME, and NJ **(Figures 3, 4, 5 and 6 Phylogenetic trees constructed by MP, ML, ME, and NJ method, respectively)** were all largely consistent. The genotype from the Philippines was used as the outgroup, and the other sequences clustered into 2 major branches: one composed of genotypes from Yunnan and Sichuan provinces, and the other including genotypes from nine other provinces. The branch containing Sichuan and Yunnan genotypes was further divided into two sub-branches, named “Sichuan branch” and “Yunnan branch”, respectively. The remaining nine provincial branches were divided into three sub-branches originating from Fujian, Guangxi, and the other seven provinces, respectively. The bootstrap values of all these branches were greater than 70, with most exceeding 90.

**Figure 3.**
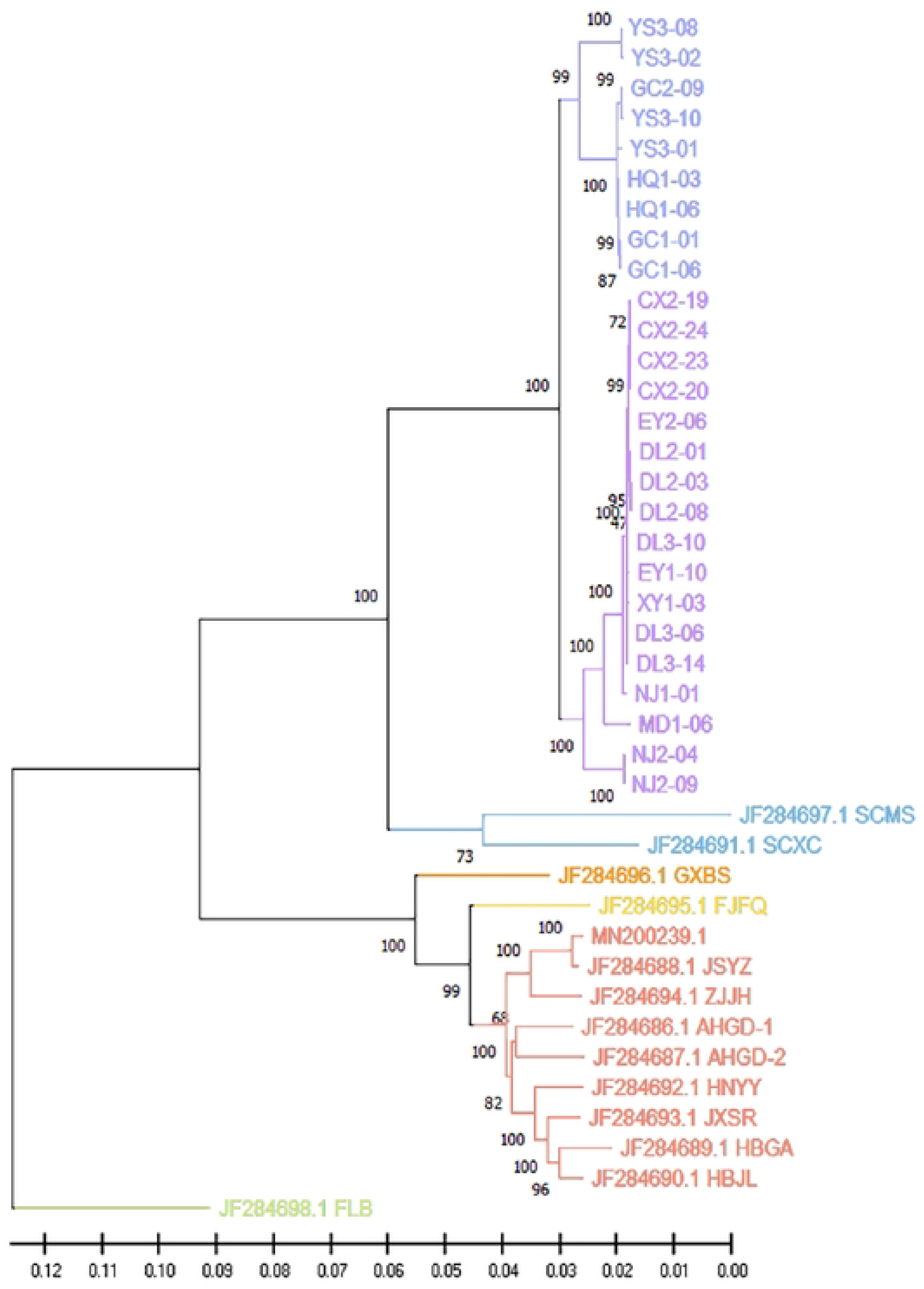
Phylogenetic tree constructed by MP method

**Figure 4.**
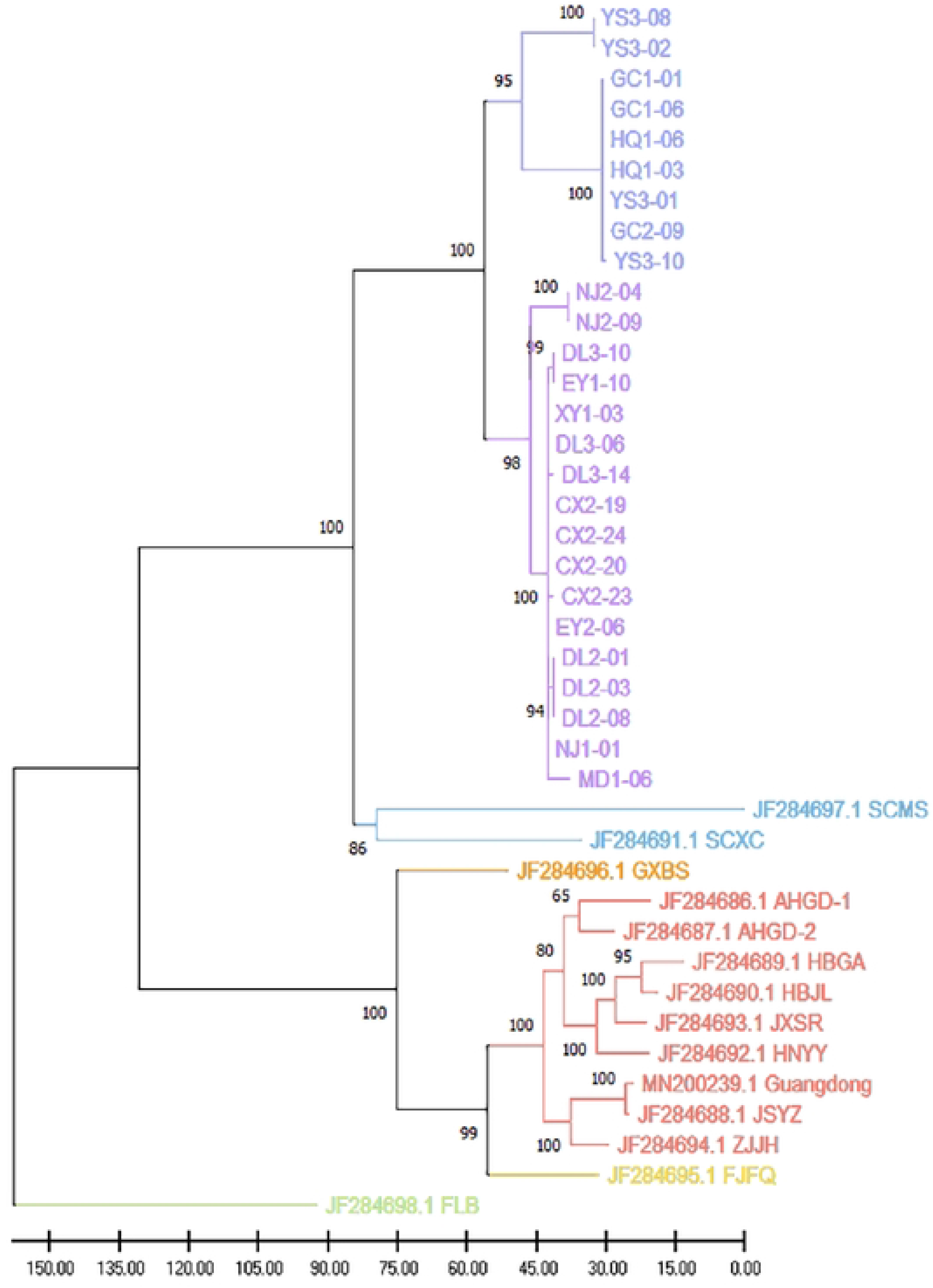
Phylogenetic tree constructed by ML method

**Figure 5.**
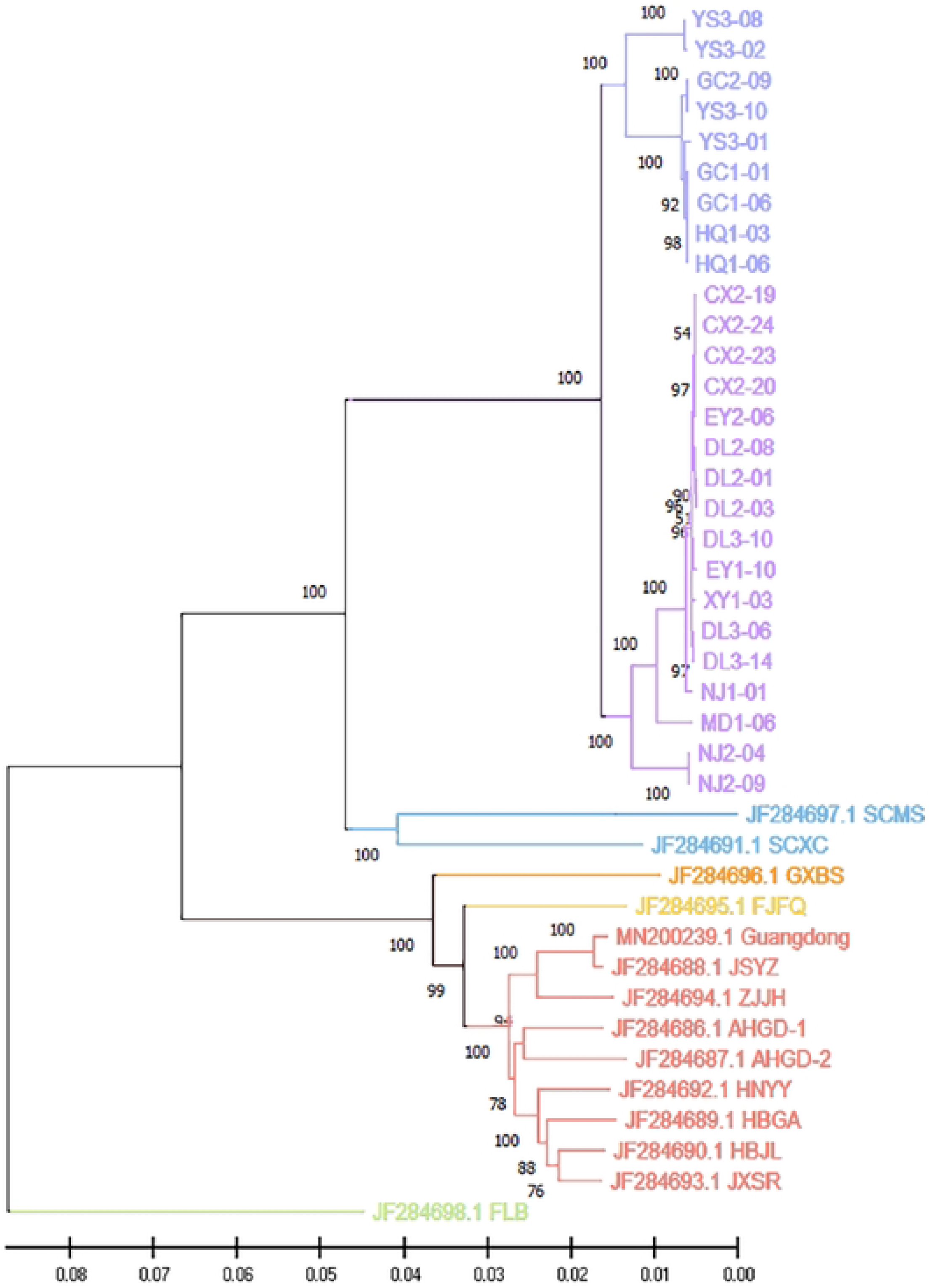
Phylogenetic tree constructed by ME method

**Figure 6.**
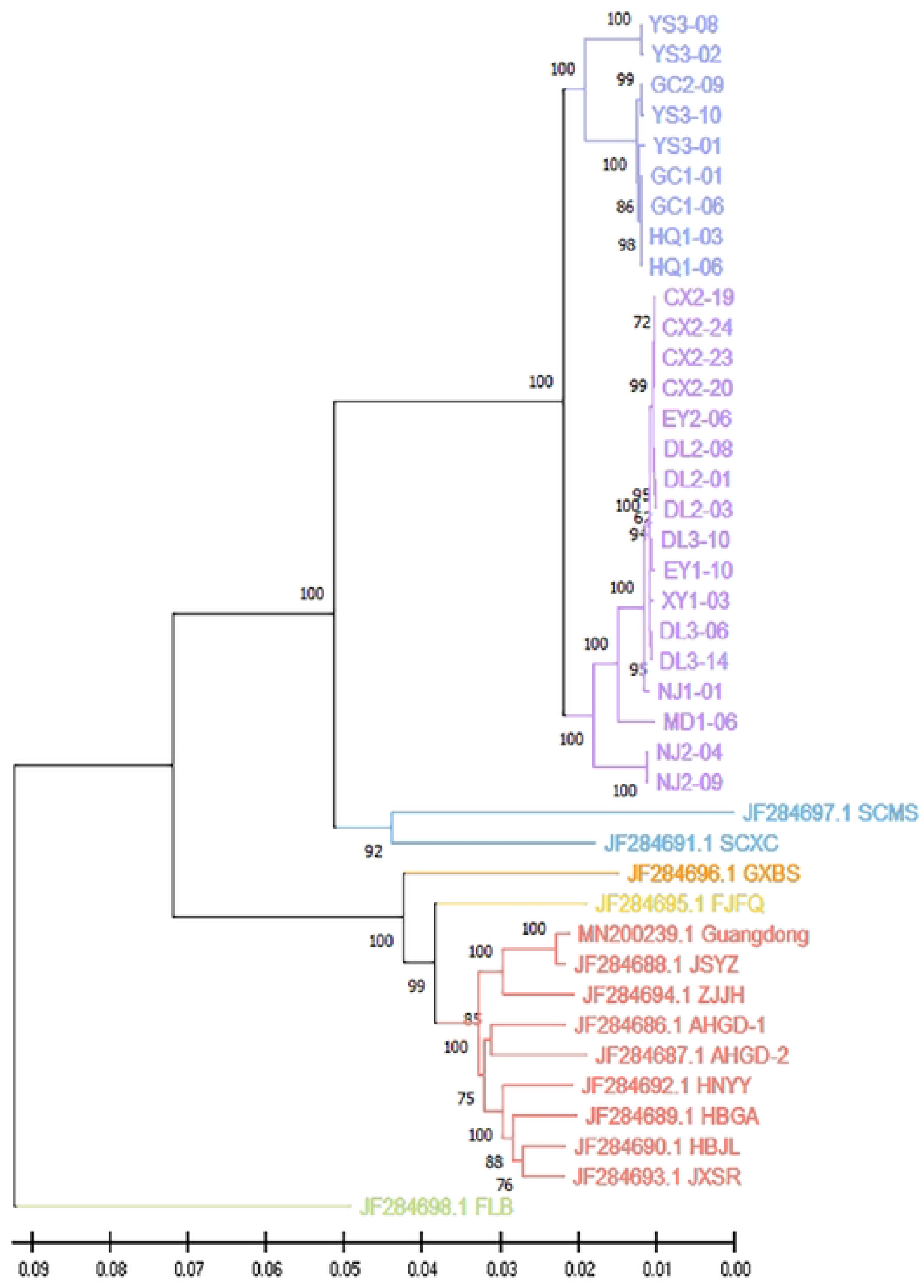
Phylogenetic tree constructed by NJ method

According to the results of phylogenetic tree, the *O. h. r.* Yunnan strain were divided into North branch and South branch. The North branch included samples from four administrative villages in three counties of Yongsheng, Gucheng, and Heqing, located in the Jinsha River Watershed in northern Yunnan Province. The South branch included samples from ten administrative villages in seven counties of Dali, Weishan, Er Yuan, Mi Du, Xiangyun, Nanjian, and Chuxiong, except for Wuxing Village in Dali City, which is located on the edge of the Jinsha River Watershed the other nine administrative villages are situated in the Lancang River Watershed, Yuan River Watershed, and Longchuan River Watershed outside the Jinsha River Watershed, in the central and southern parts of Yunnan Province. There are obvious geographic barriers between the Jinsha River Watershed and the Lancang River, Yuan River, and Longchuan River Watershed, as shown in figure1.

### 3.4 Genetic distance and homology analysis

The average genetic distance within *O. h. r.* Yunnan strain was 0.0129, while the genetic distance between the northern and southern branches was 0.0216. The average genetic distance between the *O. hupensis* of Yunnan strain and Sichuan strain was 0.0834. The average genetic distance between the *O. hupensis Gredler* (Zhejiang, Guangdong, Jiangsu, Anhui, Hunan, Hubei, Jiangxi Provinces) was 0.0216. The average genetic distance between the *O.h. robertsoni* and *O.hupensis Gredler* was 0.113. The average genetic distance among provinces of the *O.hupensis Gredler*, Fujian, Guangxi was 0.0291.

Some of the clustering information in the phylogenetic tree can be visualized in the ANI heat map (**Figure 7** **ANI heat map of mitochondrial sequence of *O. hupensis*)**, as well as the degree of homology among the samples. As shown in figure 6, the homology among the samples of *O. h. r.* Yunnan strain was above 97%, forming an orange rectangle, which contained two small red rectangles, namely “North Yunnan branch” and “South Yunnan branch”. The other subspecies of *O. hupensis*, except for the *O. h. robertsoni* and Philippine *O. hupensis*, formed a rectangle in the lower right corner.

**Figure 7.**
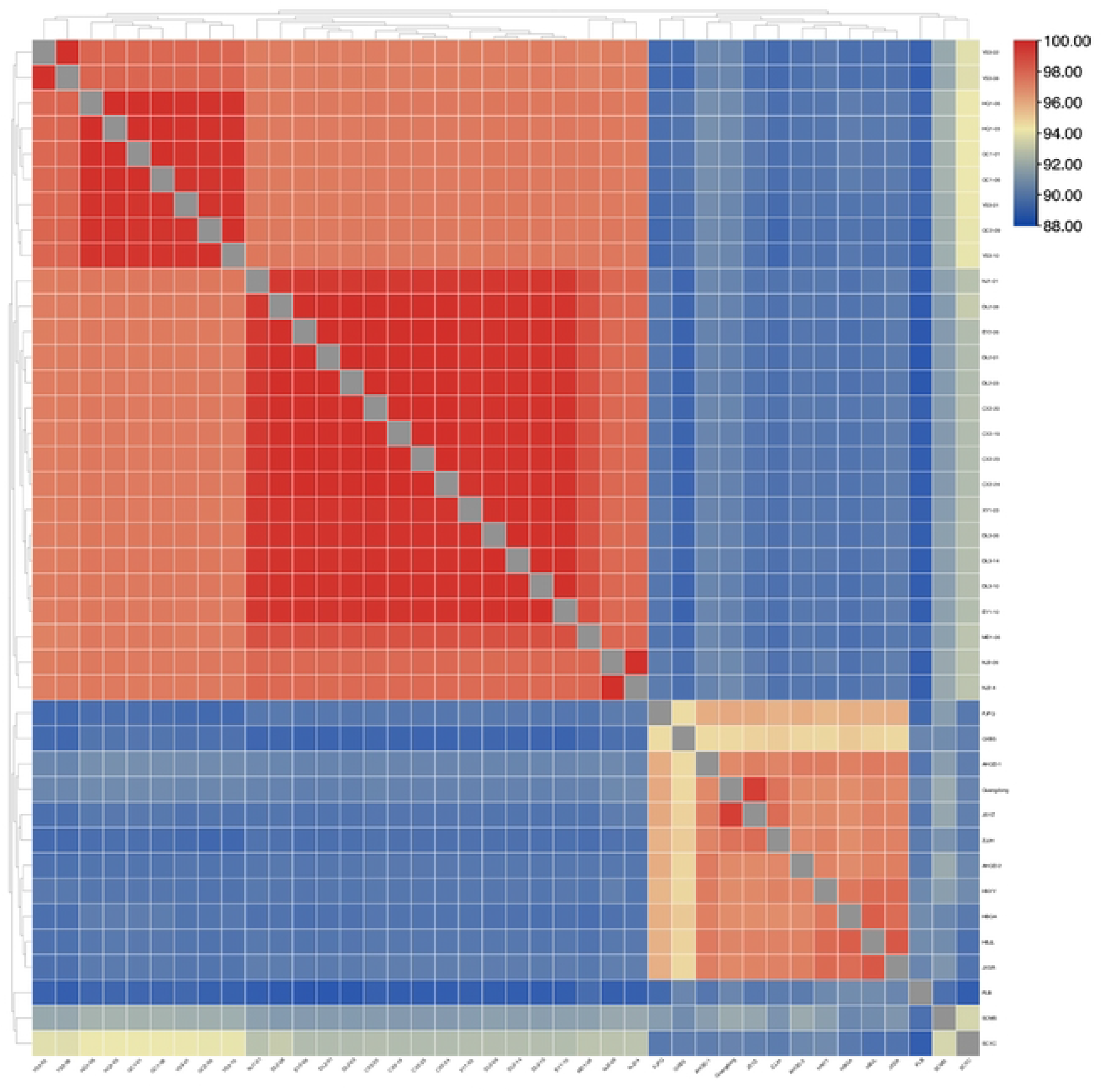
ANI heat map of mitochondrial sequence of *O. hupensis*

## 4 Discussions

In recent years, genetic sequence analysis has gained widespread utilization in the fields of phylogenetics and population genetics[25–28]. MtDNA has many advantages over nuclear genes for phylogenetic inference and classification due to its easy amplification with a large number of available conserved primers, lack of recombination, introns, and non-coding sequences, maternal inheritance, absence of recombination, which simplify the complexity of phylogenetic studies[25, 29–32]. Furthermore, mtDNA evolves rapidly, exhibits high variability within populations, and has high sensitivity for resolving closely related species[33]. Currently, mitochondrial gene fragments or mitochondrial genomes are the common markers for analyzing genetic polymorphism and genetic variation of *O. hupensis*. However, individual gene fragments may no longer provide the necessary level of identification and comprehensive phylogenetic analysis[20, 21]. While previous studies have reported the mitochondrial genome sequence of *O. hupensis* and its phylogenetic analysis, they have not conducted a systematic analysis of *O. h. r.* Yunnan strain, and the number of samples involved in the previous research on the mitochondrial genome was also limited[17]. The infection of *O. hupensis* by *S. japonicum* cercariae is an important link in the prevalence of schistosomiasis, and its susceptibility determines to some extent the extent of the disease’s spread[4]. *O. hupensis* shows different levels of genetic differentiation, leading to varying degrees of susceptibility to *S. japonicum*[34–36]. The study herein obtained 26 complete mitochondrial sequences of *O. h. r.* Yunnan strain, these sequences represent a diverse range of breeding environments characterized by different altitudes, distribution water systems, and environmental types.

In this study, the full-length sequences of the 26 mtDNA genomes ranged from 15,181 bp to 15,187 bp, with an average length of 15,185bp, in which the A+T content (67.5%) was significantly higher than the G+C content (32.5%). The mitochondrial genome encoded 37 genes, including 13 protein-coding genes, 2 rRNA genes, 22 tRNA genes and a non-coding region rich in A+T. The gene composition and distribution closely mirrored that of the mitochondrial genome of *O. hupensis Gredler*[17].

The phylogenetic trees constructed by the MP, ML, ME and NJ methods exhibited consistent topologies, and most of the branch bootstrap values of the four methods were greater than 90, indicating a high degree of reliability[37]. The clustering of Yunnan and Sichuan *O. hupensis* into one large branch, supporting the previous classification of the *O. h. robertsoni*[2, 4, 38]. In addition, the Yunnan and Sichuan populations formed two distinct branches, showing clear differentiation. It’s worth noting that the genetic distance between Yunnan and Sichuan populations was significantly greater than that observed between other subspecies of *O. hupensis*, but the genetic distance between other subspecies of *O. hupensis* was narrower, reflecting higher homology and less pronounced differentiation. These results align with those of previous studies by Han et al. and Bai, which examined the COX1 gene of *O. hupensis*, while the bootstrap values of some branches in the phylogenetic trees constructed by these two researchers were relatively small[39, 40].

Phylogenetic tree analysis showed that *O. h. r.* Yunnan strain could be subdivided into two subgroups, namely the “North Yunnan Branch” and “South Yunnan Branch”. However, when considering the ANI heat map and genetic distance, the average genetic distance within the *O. h. r.* Yunnan strain was 0.0129, indicating a high degree of homogeneity and limited differentiation. Although the *O. h. r.* Yunnan strain is divided into two small subgroups, genetic differentiation between these subgroups is not significant, as evidenced by a genetic distance of 0.0216, which is similar to the average genetic distance observed between provinces of *O. hupensis Gredler* in this study. *O. h. r.* Yunnan strain is mainly distributed in three river watersheds: the Jinsha River, Lancang River, and Yuan River. Their distribution pattern is fragmented by high mountain barriers or isolation, resulting in three distinct regions: western Yunnan, central Yunnan, and southern Yunnan. Even within the same county, the distribution areas may be characterized by small or point-like “islands” due to mountainous terrain or river barriers. Previous research has explored the presence of these “island” characteristics in *O. h. r.* Yunnan strain[22], but according to the data in this study, there isn’t a clear “island” feature. Instead, these seems to be a correlation with the distribution pattern of the river watershed. Notably, the *O. hupensis* in the “North Yunnan” subgroup exclusively occupies the Jinsha River watershed, while the remaining sampling points are distributed across the other three river watershed, with the exception of Wuxing Village in Dali City, located on the southern edge of the Jinsha River Watershed. Although three river watersheds are separated by obvious geographical barriers, samples from each watershed cluster together in the same branch of the phylogenetic tree, indicating a high degree of homogeneity and no apparent differentiation based on the distribution of river watershed. The divergence between the northern and southern lineages of Yunnan snails may result from either unique natural factor in the Jinsha River Watershed or geographic isolation due to the significant distance separating the two lineages. Further research is required to determine the exact cause of this divergence. In addition, Wuxing Village in Dali City, situated within the Jinsha River Watershed, grouped with the “South Yunnan” lineage on the phylogenetic tree. This deviation may be due to its position on the edge of the Jinsha River Watershed, making it more susceptible to gene flow with other snail populations in Dali that are geographically closer.

Furthermore, the genetic distance between the two geographical strains in Sichuan Province was 0.0704, surpassing the average genetic distance of 0.0291 observed among the other subspecies of *O. hupensis*, with the exception of *O. h. robertsoni* and the Philippine *O. hupensis*, this suggested a lower degree of homogeneity and more pronounced differentiation. However, only two geographical strains of *O. hupensis* from Sichuan province were included in this study, and further research with larger sample sizes is needed to verify the actual differentiation status of *O. hupensis* in Sichuan Province.

Major strengths of this study lie in obtaining the complete mitochondrial genome data of *O. h. r.* Yunnan strain, and we conducted a comprehensive analysis of the genetic differentiation using constructing phylogenetic trees, calculating genetic distances, and analyzing homogeneity, which ensured a high level of reliability in our results[37]. However, there is limitation to this study. Due to extensive schistosomiasis control efforts, the snails in Gejiu city, located in the southern part of Yunnan Province, have been eliminated[41]. Consequently, during *O. hupensis* sampling for this study, we were unable to collect *O. hupensis* from this region. Gejiu city was relatively isolated within the previous endemic regions of schistosomiasis in Yunnan Province, the genetic differentiation analysis of *O. hupensis* in this area was not included, which may introduce some limitations to our findings.

## 5 Conclusion

In summary, our study successfully obtained the complete mitochondrial genome of *O. h. r.* Yunnan strain and conducted a comprehensive analysis of its mitochondrial genome phylogeny. Our results indicate that *O. h. r.* Yunnan strain exhibits a higher level of genetic homology and clear north-south differentiation. Notably, *O. h. r.* Yunnan strain does not exhibit distinct “island” characteristics but instead shows a close association with watershed distribution. Taken together, the findings provide a molecular biological basis for the study of genetic evolution and resistance genes of *O. hupensis*. Additionally, it may provide useful clues for the development of snail-killing drugs, and serve as a theoretical basis and scientific evidence for the control of schistosomiasis, particularly with regard to snail control.

## Acknowledgements

We thank schistosomiasis control institutions in Yunnan Province for their valuable helps in *O. hupensis* collection.

## 6 Author contributions

Jing Song, Hongqiomg Wang, Yuwan Hao, Chunhong Du, Yi Dong conceived and designed the study. Yi Dong, Chunhong Du, Jing Song, Zongya Zhang, Jihua Zhou, Chunying Li collected samples. Yun Zhang drew sample map. Jing Song, Chunhong Du, Zongya Zhang, Meifen Shen, Chunqiong Chen, Lifang Wang, Jiayu Sun Performed the experiments. Peijun Qian, Wenya Wang, Jing Song, Hongqiomg Wang, Yi Dong analyzed, and interpreted the data. Jing Song, Hongqiomg Wang, Sizhu Li, Yi Dong drafted the manuscript. All authors critically revised the manuscript for scientific content and approved the final version of the article.

